# Bottlebrush Polymer Conjugates for Enhanced Antisense Oligonucleotide Therapy in Myotonic Dystrophy Type 1

**DOI:** 10.1101/2025.08.06.668589

**Authors:** Yao Li, Christopher Oetheimer, Yuyan Wang, Gyu Seong Heo, Jiaqi Wu, Rong Chang, Wei Zhang, Elle Schnieder, Junjie Chen, Yang Fang, Yun Wei, Keqing Nian, Hengli Zhang, Lauren Sherman, Yongjian Liu, Ke Zhang

## Abstract

Oligonucleotides are a promising genetic medicine for myotonic dystrophy type 1 (DM1), the most common adult-onset muscular dystrophy. However, poor muscle distribution of nucleic acid drugs after systemic administration has hindered drug development, and no curative treatment exists. Additionally, DM1 pathology requires drug localization to the nucleus, where pathogenic mutant RNA is trapped, posing challenges after endocytosis and endosomal escape. Here, we show that a locked nucleic acid oligonucleotide targeting mutant CUG^exp^ RNA tracts, conjugated to a bottlebrush polymer, exhibited improved muscle distribution and potent correction of DM1-associated splicing at low nanomolar doses in a DM1 mouse model. Significant improvements in myotonia, body weight, and grip strength were observed. The conjugates were well tolerated after 12 weeks of weekly intravenous dosing. These results suggest that bottlebrush polymer bioconjugates may overcome key limitations of traditional antisense drugs for muscular dystrophies, with the potential as potent, durable, and cost-efficient DM1 therapies.

**Graphical Abstract:** 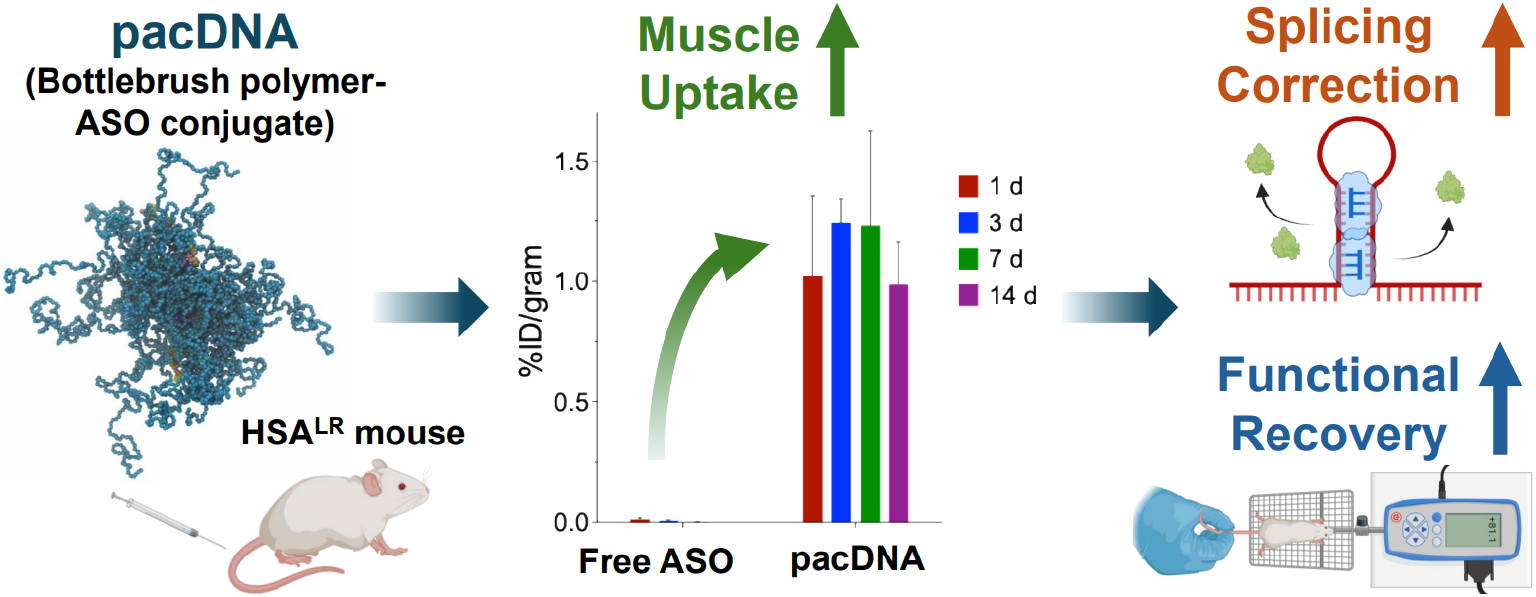

## Introduction

Myotonic dystrophy type 1 (DM1) is a multisystem muscle-wasting disease caused by an unstable CTG repeat expansion mutation in the 3’ untranslated region (UTR) of the DM1 protein kinase (DMPK) gene. DM1 affects 1 in 2,100 individuals and is the most common form of adult-onset muscular dystrophy (*1*). The autosomal dominant mutation lengthens in successive generations, leading to increased disease severity and earlier symptom onset (*2*). Pathogenesis is primarily attributed to the loss of function of muscleblind-like (MBNL) RNA-binding proteins, key regulators of pre-mRNA splicing. MBNL proteins are observed to colocalize with DMPK CUG-repeat (CUG^exp^) RNA hairpins, forming large nuclear aggregates or “foci”, leading to altered signaling pathways and gene splicing abnormalities (*3*). Loss of DMPK alone does not result in major features of DM1, and thus, transgenic mouse models with long (CTG)_n_ gene insertions (e.g., HSA-LR20b, with a (CTG)_220_ human *Acta1* transgene) have been developed to study DM1 (*4*).

Antisense oligonucleotides (ASOs) are a growing class of drugs with attractive qualities, including high specificity, customizability, and robust manufacturing. ASOs exert their therapeutic effect by hybridizing and blocking or enzymatically degrading target RNA. ASOs have gained considerable recent commercial successes, as seen in multiple exon-skipping therapeutics for Duchenne muscular dystrophy (DMD) and a ground-breaking spinal muscular atrophy (SMA) drug, highlighting the promise of ASOs for neuromuscular diseases (*5, 6*). In DM1, targeting CUG^exp^ RNA can prevent secondary structure formation and limit entrapment of MBNL proteins, leading to recovery in the alternative splicing responsible for the DM1 phenotype (*3, 7–9*). Interestingly, the length of CTG_n_ mutations has been directly related to disease severity. In preclinical studies, small interfering RNA (siRNA) and steric-blocking ASOs have been reported to achieve significant repeat RNA reduction accompanied by partial splicing correction (*8, 10–12*).

Traditional ASO efficacy in muscle is limited by rapid renal clearance, low membrane permeability, and poor biodistribution (*13–16*). Various modifications, including 2’ modifications, internucleotide linkages, and backbone replacements, have been developed to increase nuclease resistance, elevate duplex stability, and improve tropism to targeted tissues (*17, 18*). Several delivery strategies have also been employed to increase ASO muscle biodistribution, endocytosis, and nuclear trafficking, including conjugation with cationic cell-penetrating peptides (CPP) and transferrin-receptor 1 (TfR1) targeting antibodies (*12, 19*). While TfR1-targeted clinical candidates show potential in early studies, poor splicing restoration, innate/adaptive immune responses, hematopoietic toxicity, and membrane perturbation have plagued current approaches (*20–25*). To date, there remains no approved curative drug for DM1, with existing treatments focusing on symptom management (*26*).

We have shown that conjugation of oligonucleotides to branched polyethylene glycol (PEG) bottlebrush polymers can lend entropic shielding to oligonucleotides and limit their off-target interactions and immunogenicity (*27–29*). While structurally simple, these conjugates, termed pacDNA (polymer augmented conjugates of DNA), evade renal clearance due to their large size (~300 kDa, ~30 nm), improve plasma pharmacokinetics (25-fold and 19-fold increase in elimination half-life and area under the curve, respectively vs. free oligonucleotide) (*27*), and enhance biodistribution to a variety of organs and tissues through the multiple-pass effect and improved membrane adsorption (*30–34*). The key to these properties is pacDNA’s ability to shield the ASO from specific or non-specific protein interactions without interfering with target hybridization kinetics or thermodynamics (*35*). By avoiding opsonization and other DNA-protein interactions, the pacDNA eludes phagocytic clearance, limits anti-carrier adaptive immunity, and reduces all side effects that derive from these interactions, making it uniquely suitable for long-term applications. Here, we report the results of a proof-of-concept study testing a pacDNA conjugate targeting toxic CUG^exp^ RNA for the treatment of myotonic dystrophy type 1.

## Results

### PacDNA enables the targeting of CUG^exp^ nuclear foci

Bottlebrush polymers with 10 kDa PEG side chains were synthesized by ring-opening metathesis polymerization (ROMP) and purified by size exclusion chromatography as previously described (Fig. 1A, Fig. S1A) (*27*). A locked nucleic acid oligonucleotide complementary to the CUG^exp^ repeat (5’-CAGCAGCAG-3’, fully modified) was conjugated to the polymer backbone via the strained cyclooctyne copper-free click reaction (*36*). Gel electrophoresis, dynamic light scattering and aqueous gel permeation chromatography demonstrate that the pacDNA-L9 particles are uniform in size, free from aggregation, and possess a near-neutral surface charge (zeta potential ζ ~ −2.9 mV compared to ζ ~ −38.0 mV for the free ASO)(Fig. S1B,C,D) (*27, 29*).

**Fig. 1.**
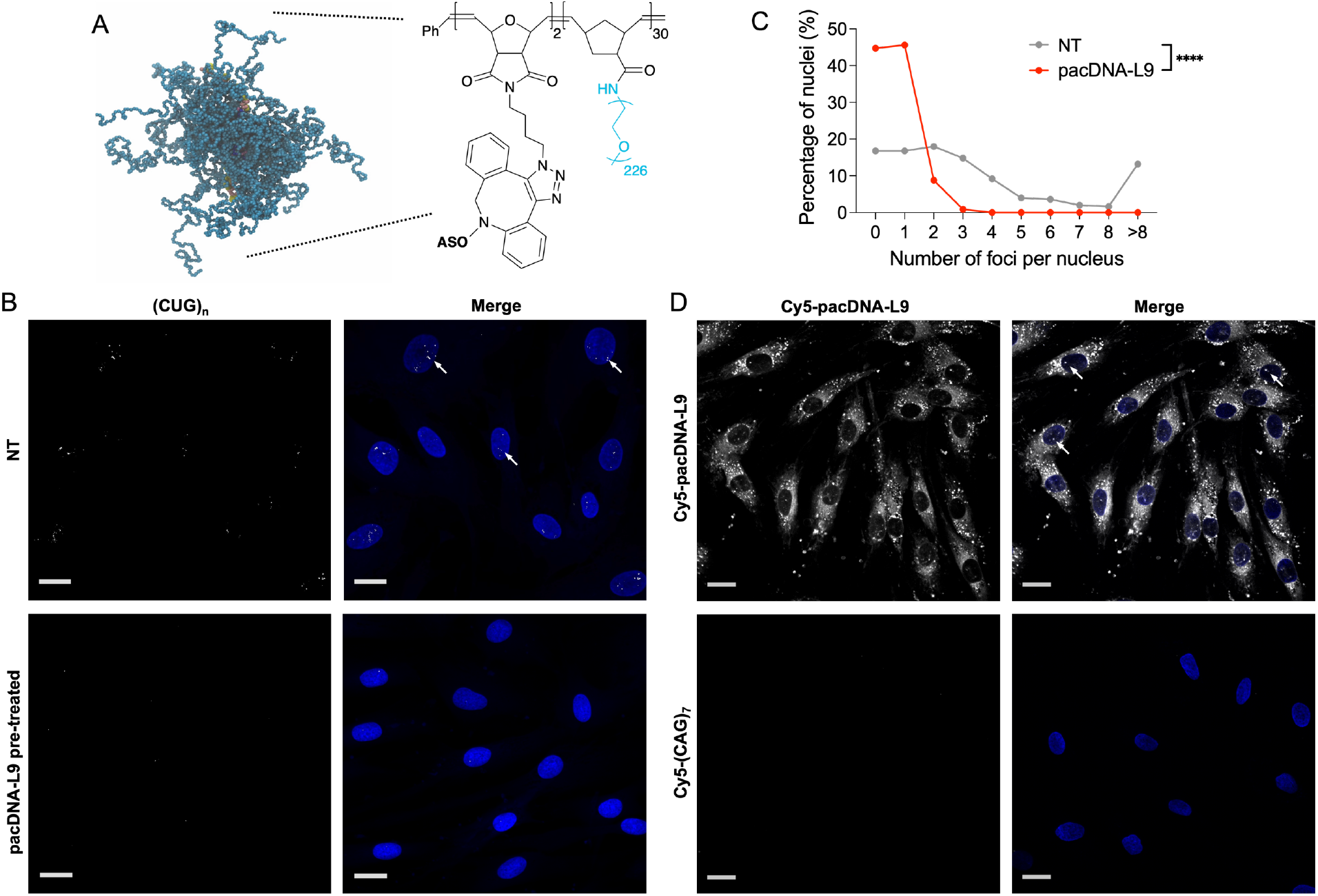
Fluorescence *in situ* hybridization in primary human DM1 fibroblasts. **(A)** Chemical structure and molecular dynamics (MD) simulation of the pacDNA in water. **(B)** Fluorescence *in situ* hybridization of primary human DM1 fibroblasts (GM03989, 2000 CTG repeats) showing CUG^exp^ foci (white, highlighted with arrows) and nuclei (blue)(upper two panels). FISH images of GM03989 cells following treatment with 1 µM pacDNA-L9 for 24 h (lower two panels) **(C)** Quantification of # of foci per nucleus in nontreated and pacDNA-L9 treated cells (250 nuclei counted per sample) **(D)** GM03989 cells treated with 1 µM Cy5-pacDNA-L9 (white) for 24 h. Engagement with nuclear foci highlighted by white arrows (upper two panels). GM03989 cells treated with 1 µM free Cy5-(CAG)_7_ (white) for 24 h (lower two panels). Scale bars: 20 µm. (****p<0.0001, nonparametric t-test.)

Fluorescence *in situ* hybridization (FISH) was performed in primary human DM1 fibroblasts with a (CTG)_2000_ repeat mutation in the DMPK gene (GM03989, Coriell Repositories). The CUG^exp^ foci characteristic of DM1 were detected as punctate patterns within the cell nuclei (Fig. 1A). When the cells were pre-treated with 1 μM pacDNA-L9 for 24 h, the number of foci detected by FISH per nucleus decreased (Fig. 1B), suggesting that the pacDNA-L9 conjugate or enzymatically liberated L9 fragments can enter the nucleus and engage with target CUG^exp^ RNA without a transfection agent. Statistically, the percentage of nuclei with no detected foci increased from 16.8% to 44.7%, with all counted nuclei in the treated group possessing three or fewer foci (Fig. 1C). When live cells were treated with fluorescently labeled Cy5-pacDNA-L9 or free Cy5-(CAG)_7_ for 24 h and imaged, the conjugate displayed improved cell and nuclear uptake relative to free ASO. The pacDNA showed nuclear accumulation in distinct positions, suggesting its ability to engage with pathogenic target foci, while the free ASO was undetectable in cell nuclei (Fig. 1D). These results indicate that pacDNA-L9 can engage with nuclear targets despite its large size.

### pacDNA enhances pharmacokinetics and biodistribution to cardiac and skeletal muscles

To evaluate pacDNA’s muscle uptake, a 14-day longitudinal biodistribution study was carried out using CD-1 mice and ^89^Zr-radiolabeled pacDNA bearing a poly(T) sequence. The probe oligonucleotide was synthesized with two desferrioxamine (DFO, siderophore-derived chelator) functionalities per molecule, which was used to chelate ^89^Zr-oxalate. Radionuclides offer better real-time, quantitative, and deep-tissue imaging capabilities compared to fluorescence-based techniques. The use of ^89^Zr, which has a relatively long t_1/2_ of 3.27 days, makes it possible to monitor the injected dose for up to two weeks. *In vitro* studies in mouse serum demonstrated >95% stability after 9 days, ensuring the accurate tracking of radiolabeled conjugates *in vivo*. CD-1 mice were i.v. injected with approximately 500 MBq of sample in 100 μL saline. At predetermined time points (24 h, 3 d, 7 d, and 14 d post-injection, n=4/time point), organs/tissues of interest were collected, weighed, and radio-counted (Fig. 2A,B). Elevated retention of pacDNA conjugate was observed in blood pool organs (blood, heart, lung), reasonable uptake in mononuclear phagocyte systems (liver, spleen), and considerable uptake in other tissues including the abdominal and thigh muscles (~1% injected dose [ID]/g) and skin (~4% ID/g). Approximately 40% of the injected pacDNA dose was cleared via the urine and feces by day 14. Importantly, given that muscle tissues constitute approximately 40% of the mass of the animal (37), at 1% ID/g, pacDNA enables ~13% of total injected dose to localize to muscles, representing a 10^2^ to 10^4^-fold increase over the free oligonucleotide depending on the time point (Fig. 2C). These muscle uptake results also compare favorably with TfR1-targeted antibodies, showing 2 to 7-fold higher muscle concentration (*38*). Muscle selectivity, as defined by the ratio of C_max_ in the dose-limiting organ (kidney) vs. muscle, also improves significantly (from ~1680 for free oligonucleotide to 20.4). Notably, the pacDNA conjugate also showed increased accumulation in the gallbladder, pancreas, and brain compared with free ASO. Positron emission tomography (PET)/computed tomography (CT) scans were carried out on day 1 and day 7 before the sacrifice of the mice (Fig. 2D). PET/CT imaging shows extended blood retention and significant tissue uptake, while the free oligonucleotide suffered from rapid renal clearance with little accumulation in non-kidney tissues. A similar biodistribution profile was observed for pacDNA-L9 in HSA^LR^ transgenic mice, demonstrating comparable accumulation in skeletal and cardiac muscles (~1% ID/g) as CD-1 mice (Fig. S2). These data affirm that pacDNA achieves robust and reproducible muscle targeting across both healthy and disease-relevant models using orthogonal tracking strategies. Collectively, these data indicate that the pacDNA is a long-circulating, long-retention vector viable for systemic delivery to address muscular disorders.

**Fig. 2.**
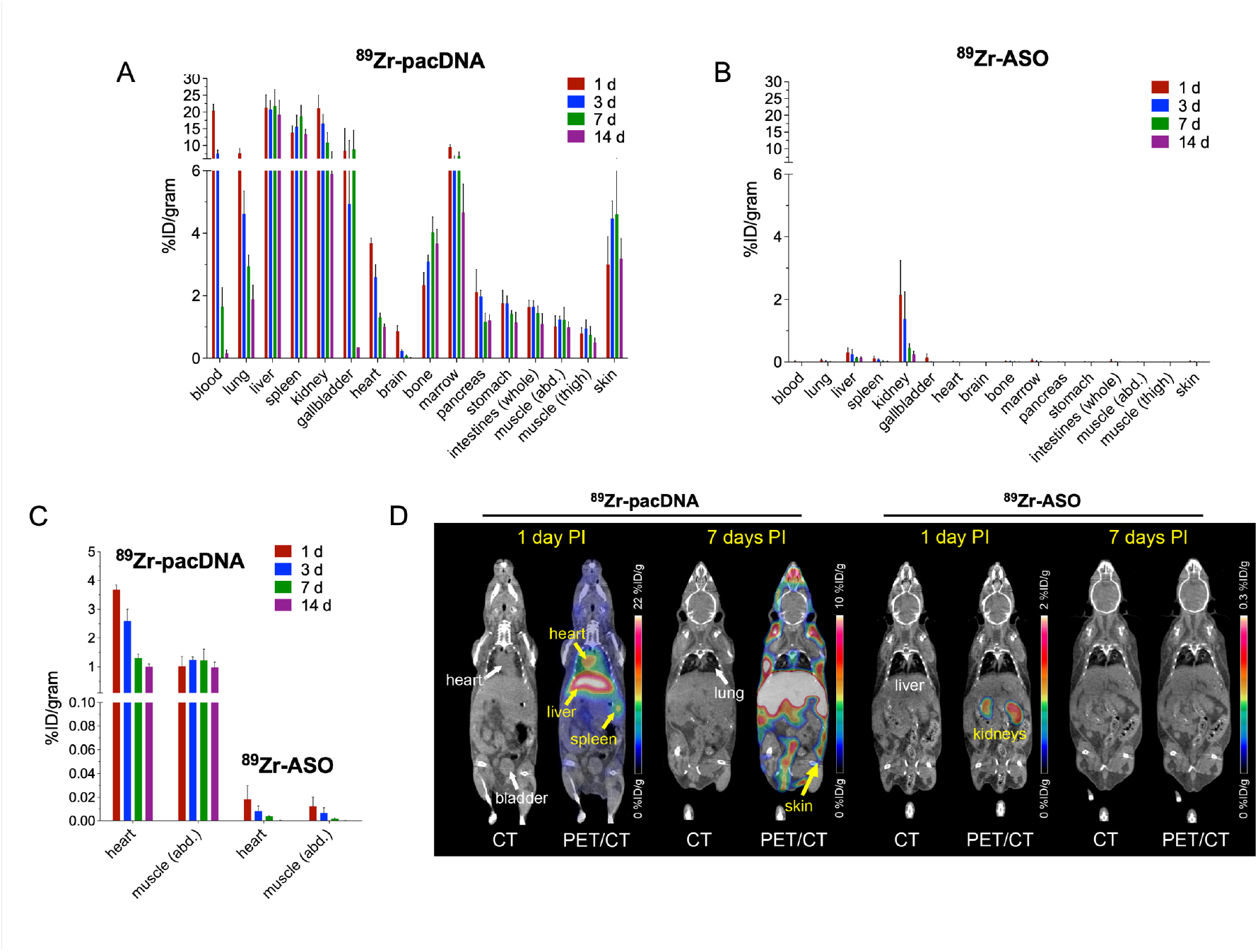
Biodistribution of radiolabeled pacDNA and free ASO. **(A, B)** Ex vivo biodistribution of ^89^Zr-labeled pacDNA and ^89^Zr-ASO in various tissues and organs measured over 14 days (n=4/time point) in CD-1 mice. The y axis shows the percent of injected dose per gram of tissue (%ID/g). **(C)** %ID/g comparison of heart and muscle in of ^89^Zr-pacDNA (left) and ^89^Zr-ASO injected mice (right). **(D)** Positron emission tomography (PET) scans of ^89^Zr-pacDNA (left) and ^89^Zr-ASO injected mice (right) at 1 d and 7 d post-injection. Scale bars represent %ID/g. Error bars indicate ±s.d.

### pacDNA-L9 corrects DM1-associated alternative splicing in HSA^LR^ mice

HSA^LR^ mice (8 weeks old) were injected i.v. by tail vein with 5.3 mg/kg-42.4 mg/kg (ASO basis) of pacDNA-L9, pacDNA with a scrambled sequence (pacDNA-Scr), free L9 control, or free bottlebrush polymer with no attached ASO. Total cellular RNA from mouse quadriceps was sequenced to evaluate gene expression and RNA splicing patterns. Among the many genes affected in DM1 are several encoding membrane ion channel proteins. Deficiencies in these proteins disrupt normal muscle ion gradients and electrophysiology, leading to hallmark physical manifestations of DM1, such as myotonia, myopathy, and arrhythmia (39-43). We measured the exon inclusion, or percent spliced-in (PSI), across a panel of 21 exonic gene fragments reported to experience mis-splicing in DM1 muscles (Fig. 3) (*44–49*). Many of these 21 genes have been identified as direct targets of MBNL splicing proteins and are therefore valuable biomarkers of MBNL protein activity in muscle. Splicing dysregulation improved two weeks after pacDNA treatment in a dose-dependent manner. Across the entire panel, the composite scaled PSI (cPSI) decreased with escalating dosing (Fig. 4A). The 5.3 mg/kg, 21.2 mg/kg, and 42.4 mg/kg cohorts exhibited average PSI corrections of 31.3%, 48.5%, and 65.5%, respectively. Notably, several key transcripts associated with specific DM1 phenotypes (e.g., *Clcn1, Insr, Cacna1s, Cacna2d1*)(*41, 50*) displayed strong splicing correction nearly back to wildtype (WT) levels. Contrastingly, in mice injected with free L9 control (5.3 mg/kg), the average PSI correction across this panel was 11.4%, with many of these genes displaying no correction or worsened spliceopathy (Fig. S3). Free brush polymer or pacDNA-Scr controls failed to correct alternative splicing in key DM1-related genes (Fig. S4). Additionally, treatment with pacDNA-L9 corrected a significant fraction of global alternative splicing. Out of 207 skipped exon (SE) events significantly altered (FDR < 0.01) in non-treated HSA^LR^ mice vs. wildtype animals, treatment with pacDNA-L9 (42.4 mg/kg) corrected the PSI of 50 events by 50-100%, 81 by 20-50%, and 39 by less than 20% (Fig. 4B). Only 37 SE events were not corrected.

**Fig. 3.**
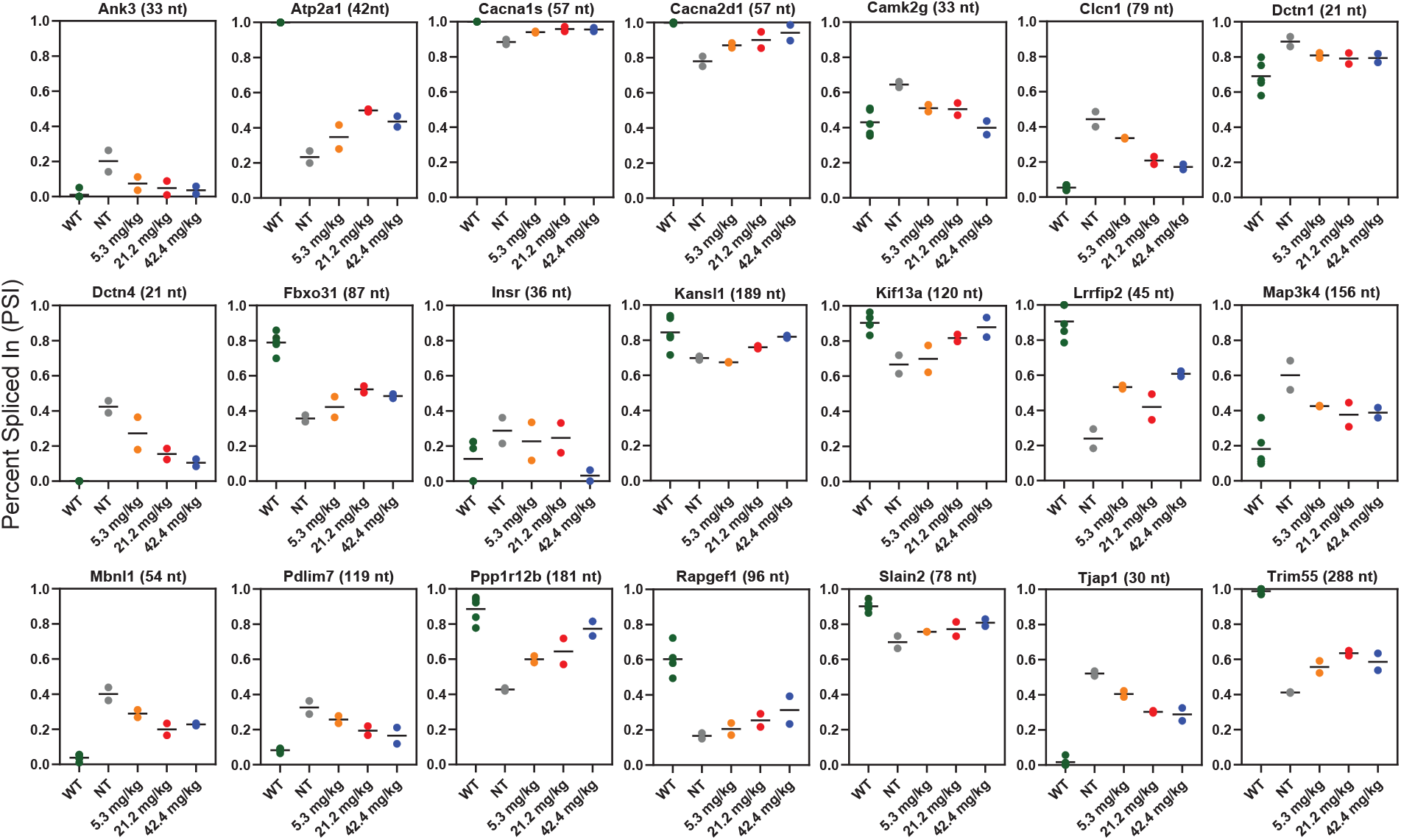
DM1 Alternative Splicing Panel. Percent spliced-in (PSI) of 21 DM1-associated splicing events in wildtype (n=5), nontreated HSA^LR^ (n=2), or pacDNA-L9 5.3 mg/kg (n=2), 21.2 mg/kg (n=2), or 42.4 mg/kg (n=2) treated groups.

**Fig. 4.**
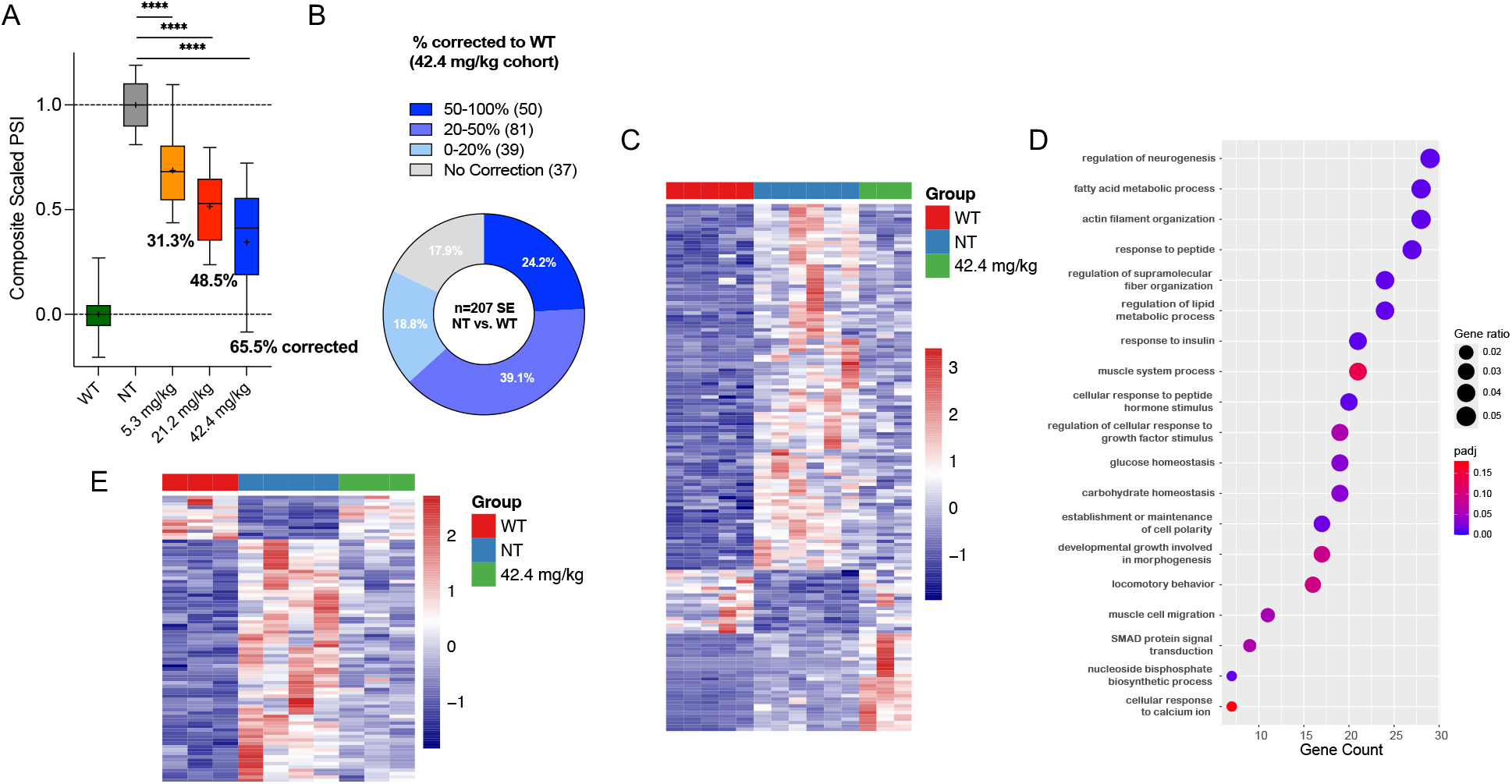
Transcriptomic alterations following pacDNA-L9 treatment. **(A)** Composite scaled PSI across the 21-gene fragment panel (Fig. 3) of DM1-associated splice events in the WT, NT, or pacDNA-L9 treated cohorts (One-way ANOVA with Tukey’s multiple comparison test, ****p < 0.0001). Percent values under each group represent the mean percent corrected to wildtype PSI levels. Tukey box-and-whisker plots indicating median (line), mean (+), interquartile range (IQR; box), and 10th and 90th percentiles. **(B)** % corrected to wildtype PSI of 207 significant skipped exon (SE) events in nontreated HSALR mice by 42.4 mg/kg pacDNA-L9 treatment **(C)** Top 25% most recovered genes following 42.4 mg/kg pacDNA-L9 treatment, visualized by heatmap. **(D)** Biological processes identified by gene set enrichment analysis (GSEA) of genes corrected (padj > 0.05 vs. WT) by 42.4 mg/kg pacDNA-L9 treatment. **(E)** Top 25% most recovered exon bins following 42.4 mg/kg pacDNA-L9 treatment, visualized by heatmap.

### pacDNA-L9 shifts the global transcriptome towards wildtype at both gene expression and splicing levels

To examine the overall effect of pacDNA-L9 treatment at the transcriptomic level, we performed bioinformatic analysis of both gene expression and alternative splicing. From all differentially expressed genes between WT and NT samples (padj < 0.05, |log2FoldChange| > 1), the top 25% most recovered genes following 42.4 mg/kg pacDNA-L9 treatment were visualized by heatmap. The calculation of a recovery score is described in the Methods. The treatment group (green) exhibited an intermediate recovery pattern, closer to the WT group (red) than to the NT group (blue), suggesting significant restoration of gene expression (Fig. 4C). Similarly, transcriptomic analysis of exon bins (normalized counts) revealed global correction of alternative splicing mis-regulation following treatment (Fig. 4E). Gene set enrichment analysis (GSEA) of significantly corrected genes (padj > 0.05 vs. WT) revealed functional signatures associated with the restored gene expression (Fig. 4D). Among the top corrected biological processes are fatty acid/lipid metabolic process, glucose homeostasis, and response to insulin, indicating a recovery in metabolism- and energy storage/conversion-related gene expression. Additionally, categories such as muscle system process, actin filament organization, muscle cell migration, and growth factor and morphogenesis suggest restoration of healthy muscle function and growth (*44, 49, 51, 52*). Gene set variation analysis (GSVA) of the 100 lowest score (most downregulated) gene sets in nontreated HSA^LR^ samples compared to wildtype highlighted the profound recovery in disease-related gene dysregulation following pacDNA-L9 treatment even in the low-dose cohort (Fig. S5, A, B). It has been shown that CUG^exp^ RNA can be sliced by Dicer into small repeated RNAs and used as templates for RNA interference (RNAi) of endogenous mRNA transcripts in the cytoplasm, contributing to global alterations of the DM1 transcriptome (*51, 53*). Due to the pacDNA’s high muscle concentrations and cell uptake, even the lowest dose (5.3 mg/kg) may be adequate for mitigating these cytoplasmic RNAi effects. Since the (CAG)_3_ ASO on the pacDNA-L9 has the potential to bind other endogenous (CUG)_n_-containing transcripts, we examined the expression levels of 22 genes with varying (2-25) CTG repeats (Table S1) (*10, 11, 54*). None of these genes were significantly up or downregulated after pacDNA-L9 treatment relative to non-treated controls, confirming the absence of off-target modulation of short (CUG)_n_-containing transcripts at these doses (Fig. S5C).

### pacDNA-L9 alleviates myotonia and improves muscle strength in HSA^LR^ mice

Next, we evaluated the phenotypic response of the pacDNA-L9 treatment in HSA^LR^ mice. DM1 in humans is characterized by progressive muscle wasting and weakness, alongside myotonia, which refers to the delayed relaxation of muscles after contraction. Myotonia is a key feature in both DM1 patients and the HSA^LR^ mouse model and contributes significantly to physical impairment. HSA^LR^ mice (8 weeks old) were administered i.v. with pacDNA-L9 at 10.6 mg/kg/day for four consecutive days, followed by weekly (10.6 mg/kg/week) dosing from week 2 to week 12. A blinded hindlimb pinch test was performed each week to evaluate myotonia (delayed muscle relaxation). Significant improvements in myotonia severity and recovery time were observed starting in the third week of treatment (p < 0.01), with maximal therapeutic effect achieved by the seventh week. Myotonia scores for the treated group remained consistently lower than those for the non-treated group (p < 0.001 from weeks 7 to 12, Fig. 5A). Additionally, mice in the treatment group showed a consistent and notable increase in body weight compared to the non-treated group, with the male group showing a 14.6% increase and the female group showing a 9.1% increase over non-treated mice of the same age at week 12 (Fig. 5B). To further assess the functional impact of pacDNA-L9 treatment, we conducted a grip strength test to evaluate muscle strength in HSA^LR^ mice. At week 12, the treated mice were significantly stronger than their untreated littermates (p < 0.001), close to wildtype levels (Fig. 5C). These findings suggest that pacDNA-L9 effectively improves muscle strength and alleviates myotonia following treatment. Further, repeated dosing does not lose efficacy over time, suggesting little to no anti-drug adaptive immunity and accelerated clearance.

**Fig. 5.**
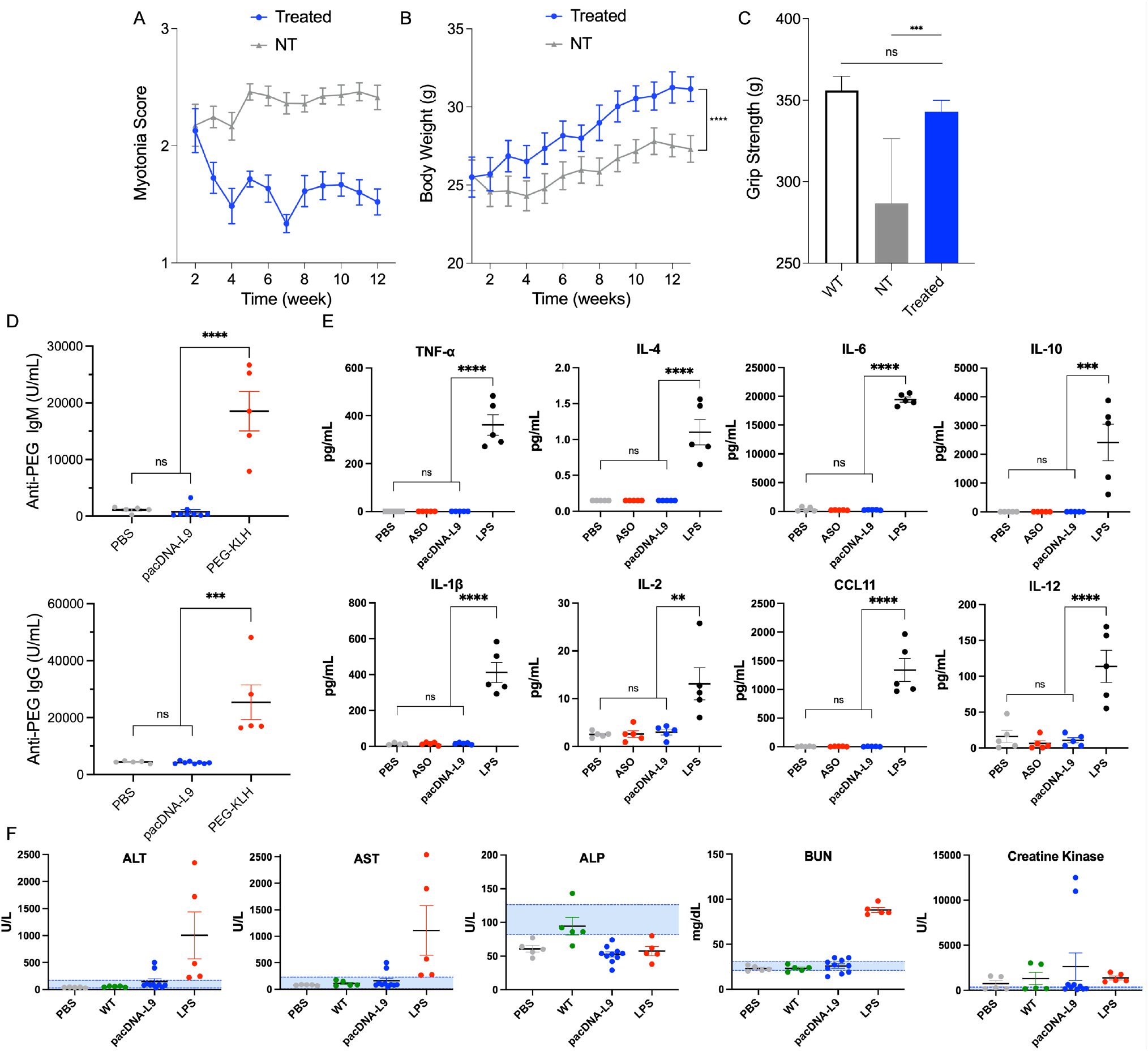
Functional effects of pacDNA-L9 treatment in HSA^LR^ mice and safety evaluation. HSA^LR^ mice received intravenous injections of pacDNA-L9 (10.6 mg/kg/dose) over 12 weeks **(A)** Myotonia scores measured weekly. The response to each pinch was classified as severe myotonia (>1 s, 3), myotonia (0.5-1 s, 2), quick recovery myotonia (<0.5 s, 1) myotonia, single leg myotonia (0.5), or no myotonia (0). **(B)** Body weight measurements were recorded throughout the administration period. **(C)** Week 12 grip strength analysis demonstrates improved muscle strength in the treated group compared to the non-treated group. The mean and SEM (n=10 per group) in wildtype (WT), nontreated HSA^LR^ (NT), and treated 10.6 mg/kg/dose HSA^LR^ are displayed (**p < 0.01, ***p < 0.001, ****p < 0.0001 by one-way ANOVA followed by Tukey multiple comparison correction). **(D)** Anti-PEG IgM and IgG antibody generation evaluated after 16 doses (10.6 mg/kg) of pacDNA-L9 by ELISA (n=8). PBS (n=5) was used as the negative control and PEG-KLH (n=5) as the positive control. **(E)** Cytokine and chemokine levels in HSA^LR^ mice serum 4 h after i.v. injection of PBS, free L9-ASO, pacDNA-L9 (10.6 mg/kg), or LPS (2 mg/kg) **(F)** Liver and kidney function biomarker levels in HSA^LR^ mice serum after i.v. injection of PBS, pacDNA-L9 (10.6 mg/kg), or LPS (2 mg/kg), or in FVB/N wildtype (WT) mice. The blue shaded area represents the healthy reference range in FVB/N mice. (**p < 0.01, ***p<0.001, ****p<0.0001, one-way ANOVA). Error bars indicate ±s.e.m.

Macromolecular therapeutics can induce adaptive immune responses that lead to loss of drug activity for subsequent doses. To measure this effect, anti-drug immunogenicity after 16 doses of pacDNA-L9 was evaluated for anti-PEG IgM and IgG response. In serum samples collected 1 week after the final injection, minimal anti-PEG IgM and IgG antibodies were detected, while the PEG-keyhole limpet hemocyanin (KLH) positive control group showed significant levels of both IgM and IgG (Fig. 5D). To assess acute immune system activation, a panel of cytokines and chemokines were measured using HSA^LR^ mouse serum collected 4 h after the last injection. For positive control, 2 mg/kg of lipopolysaccharide (LPS) was given to mice, which induced an acute immune response. No significant elevations in cytokine/chemokine levels were seen between pacDNA-L9-treated and non-treated mice, whereas the LPS-treat group exhibited significantly elevated levels in all 16 tested cytokines/chemokines, indicating that pacDNA-L9 does not trigger acute inflammatory responses (Fig 5E, Fig. S6). Liver function markers, including alanine aminotransferase (ALT), alkaline phosphatase (ALP), aspartate aminotransferase (AST), albumin, total bilirubin (TBIL), and total protein (TP), as well as renal function markers, such as blood urea nitrogen (BUN) and creatinine, remained within normal ranges, indicating that pacDNA-L9 treatment does not cause hepatic or renal dysfunction (Fig. 5F, Fig. S7). Taken together, pacDNA-L9 shows excellent tolerance in mice, with no evidence of significant immune system activation, toxicity, or anti-drug adaptive immunity throughout 12 weeks of repeated dosing.

## Discussion

The challenge of muscular delivery of oligonucleotides was highlighted by the discontinuation of the development of *baliforsen* (IONIS 598769), the first ASO therapy for DM1, in 2017, citing insufficient drug concentration in muscle after subcutaneous injection (*55*). Besides, local intramuscular injections are impractical, considering the systemic pathology of the disease. Transferrin receptor-targeted approaches have recently gained popularity for muscle delivery, and Avidity Biosciences and Dyne Therapeutics are in clinical trials testing a transferrin receptor 1-targeted mAb-siRNA conjugate (*del-desiran*) and a Fab-ASO conjugate (Dyne-101) for degradation of mutant DMPK mRNA (*24, 25, 56*). However, muscle biopsies from patients given *del-desiran* and Dyne-101 displayed 12% and 19% maximal correction of mis-splicing in their respective panels, indicating additional room for improvement. Further, the complex antibody/Fab conjugate may present challenges in manufacturing, considering the stability of the protein component during and after conjugation. In addition, conjugate stability may lead to difficulties in transport, storage, and handling at the clinic.

A second approach to the delivery problem is the use of polycationic substances such as lipids, polymers, or peptides, which improve per-molecule activity by enhancing cell uptake and endosomal escape (*12, 57*). While effective *in vitro* and promising in murine and non-human primate models, polycationic substances pose toxicity concerns, especially at high doses, due to nonspecific lytic activity, unwanted immune system activation, and organ damage (*58, 59*). Sarepta has recently terminated its DMD program, SRP-5051, a cationic peptide-phosphorodiamidate morpholino oligomer (PPMO) conjugate. It is thought that cationic conjugates will likely have a narrow therapeutic window (*60*). Clinical results of DM1 programs cell-penetrating peptide conjugates, such as VX-670 and PGN-EDODM1, remain to be seen.

In contrast to the existing oligonucleotide delivery modalities, the pacDNA construct is novel in both chemistry and mechanism. It achieves comparable potency as the state of the art for DM1 despite having a yet unoptimized ASO (*56, 61, 62*). After i.v. injection, the superior blood retention properties of the pacDNA facilitate the distribution of the ASO to skeletal/cardiac muscles, yielding higher muscle concentrations than Ab conjugates (by 2 to 7-fold) and peptide conjugates (>100-fold)(*38, 63*). While the improved plasma retention also leads to higher drug levels in the kidney and the liver, muscle selectivity (defined as muscle concentration relative to dose-limiting organ, e.g., kidney) also improved significantly compared to free ASO. Importantly, 12 weeks of repeated dosing did not result in any detectable toxic and immunogenic side effects. Blood biochemistry markers remained within normal range, and no cytokine response related to innate and adaptive immunity was observed. Further, the treatment did not result in anti-drug antibodies (ADA), as measured by ELISA for anti-PEG IgG and IgM. These results are consistent with our previous studies (for non-DM1 targets), showing a clean safety profile and non-immunogenicity in both murine and non-human primate models (*27, 64*). Unlike antibodies and antibody-drug conjugates, the pacDNA is stable at room temperature for at least one month and can be repeatedly lyophilized and reconstituted, providing simplicity in storage and transport and convenience at the clinic. Owing to strong safety characteristics, elevated muscle bioavailability, and a favorable drug profile, the pacDNA may be able to more reliably exceed therapeutic thresholds of splicing correction compared with current modalities (*24, 25, 46*).

As the first version of a readily adaptable platform, the pacDNA has the potential for iterative optimization via polymer backbone modifications (*30*), linker chemistries (*65*), the attachment of ligands that can increase cell targeting, nuclear uptake, or endosomal escape (*66–68*), and/or ASO optimization. This adaptability will allow the pacDNA structure to evolve alongside our understanding of what drives potency. This study serves as a basis for this fundamentally new oligonucleotide delivery modality for the treatment of DM1, opening the possibility for further development in other neuromuscular diseases such as myotonic dystrophy type 2 (DM2), DMD, and facioscapulohumeral muscular dystrophy (FSHD).

## Materials and Methods

### Experimental Design

This study evaluated the delivery efficiency, therapeutic efficacy, and safety of bottlebrush polymer–conjugated locked nucleic acid antisense oligonucleotides (pacDNA) for the treatment of myotonic dystrophy type 1 (DM1). The HSA^LR^ (HSA-LR20b) transgenic mouse model was selected for its well-characterized DM1-like features, including CUGexp RNA nuclear foci, splicing misregulation, myotonia, and muscle weakness.

To assess the delivery performance of pacDNA, we first examined its biodistribution in skeletal muscle of treated mice, and its nuclear localization in DM1 patient-derived cells, to confirm both tissue uptake and intracellular target engagement. To evaluate dose-dependent efficacy, HSA^LR^ mice were treated intravenously with pacDNA at 5.3, 21.2, or 42.4 mg/kg. To assess durability and long-term tolerability, a separate cohort received weekly intravenous injections for 12 consecutive weeks, starting at 8 weeks of age.

Therapeutic outcomes were assessed by RNA sequencing to quantify transcriptome-wide splicing correction and gene expression changes. Functional improvement was evaluated by pinch-induced myotonia scoring, grip strength testing, and body weight monitoring. Safety was evaluated through cytokine and chemokine profiling, anti-PEG IgM/IgG quantification, and serum biomarkers of hepatic and renal function.

### Materials and reagents

Methoxy PEG Amine, HCl Salt (A3052) was purchased from Jenkem Technology USA. Phosphoramidites and supplies for DNA synthesis were obtained from Glen Research Co., USA. All other materials were obtained from MilliporeSigma, Fisher Scientific Inc., USA, or VWR International LLC., USA, and were used as received unless otherwise indicated.

### Instrumentations

Reversed-phase high-performance liquid chromatography (RP-HPLC) was performed on a Waters (Waters Co., MA, USA) Breeze 2 HPLC system coupled to a Symmetry® C18 3.5 μm, 4.6×75 mm reversed-phase column and a 2998 PDA detector, using TEAA buffer (0.1M) and HPLC-grade acetonitrile as mobile phases. Aqueous GPC measurements were performed on a Waters Breeze 2 GPC system equipped with Ultrahydrogel™ 1000, Ultrahydrogel™ 500 and Ultrahydrogel™ 250, 7.8Å×300 mm column and a 2998 PDA detector (Waters Co., MA, USA). 0.1 M sodium nitrate (pH 7.0) was used as the eluent running at a flow rate of 0.8 mL/min.

*N,N*-dimethylformamide (DMF) GPC was performed on a TOSOH EcoSEC HLC-8320 GPC system (Tokyo, Japan) equipped with a TSKGel GMHHR-H, 7.8×300 mm column, and RI/UV-Vis detectors. HPLC-grade DMF with 0.05 M LiBr was used as the mobile phase, with samples run at a 0.5 mL/min flow rate. GPC calibration was performed with a polyethylene glycol/polyethylene oxide Readycal set (Sigma Aldrich, USA).

### Cell culture and animals

Human DM1 patient-derived fibroblasts with a ~2000 CTG repeat mutation in the DMPK gene and healthy control fibroblasts were obtained from the NIGMS Human Genetic Cell Repository at the Coriell Institute for Medical Research: [GM03989, GM07492]. Cells were passaged and cultured in Eagle’s Minimum Essential Medium (EMEM) with 10% fetal bovine serum (FBS), 1% antibiotic antimycotic, and 1% non-essential amino acids solution in 5% CO2 at 37° C. All cell counting was performed using Trypan Blue and a hematocytometer instrument. Cells were frozen for storage in 90% FBS with 10% DMSO (1 mL).

Homozygous HSALR20b (HSA^LR^) transgenic mice (human skeletal actin long repeat line 20b, Strain #032031) and FVB/N (Strain #001800) were purchased from The Jackson Laboratory, USA (*4*). Animal experiments were conducted at Northeastern University and carried out in accordance with approved Institutional Animal Care and Use Committee guidelines (protocol number: 22-0309). Adult mice aged ~8 weeks were injected via tail vein with pacDNA, free DNA, or scramble/brush polymer controls dissolved in 200 µL of sterile phosphate buffered saline (PBS). Animals were humanely sacrificed by CO_2_ at indicated time points for tissue collection. Blood for serum analysis assays was collected from the submandibular vein at indicated time points.

### Antisense Oligonucleotides

Custom DNA oligonucleotides were synthesized on a Dr. Oligo 48 DNA synthesizer (Biolytic Lab Performance, Inc., Fremont, CA). Short-length single-stranded DNA with a sequence of 5’-CAGCAGCAG-3’ of locked nucleic acid (LNA) modified bases was made, and dT-DBCO was attached at the 5’ end for downstream “click” reactions. All-LNA scramble control oligonucleotides (pacDNA-Scr, 5’-CGCACAAGG-3’) were synthesized in parallel. A fluorescent probe with a sequence of 5’-(Cy-3)-C**A**GC**A**GC**A**GCAG**C**AG**C**AG**C**A-3’ (bold=LNA modification) was synthesized for fluorescence *in situ* hybridization assays (*69*). All DNA was synthesized using 1 micromole controlled pore glass (CPG) dT columns, or 200 nanomole Cy-3 CPG columns.

Oligonucleotides were cleaved from the CPG solid support and deprotected in aqueous ammonium hydroxide solution (28-30% NH_3_ basis) at room temperature for 16 h. Free oligonucleotides were purified by RP-HPLC. If necessary, dimethoxytrityl (DMT) protecting groups were removed by treatment with 20 % acetic acid for 1 h followed by 3x ethyl acetate extraction. Purified DNA was quantified on a NanoDrop® ND-1000 UV-Vis Spectrophotometer and lyophilized for storage at −20° C.

### Synthesis and purification of pacDNA conjugates

Norbornenyl bromide was synthesized using a two-step synthesis involving maleimide (1 equiv.), furan (1 equiv), 1,4-dibromobutane (4 equiv.), and K_2_CO_3_ (5 equiv.) as previously described (*27*). 5-norbornene-2-acetic acid succinimidyl ester (norbornenyl NHS ester) was synthesized via a reaction of exo-5-norbornene carboxylic acid (MilliporeSigma, 1 equiv.) and N-hydroxysuccinimide (Thermofisher, 1.4 equiv.) in the presence of 1-ethyl-3-(3-dimethylaminopropyl)carbodiimide (EDCI) (MilliporeSigma, 1.3 equiv.) as a carboxyl activating agent in dichloromethane (DCM) for 16 h at room temperature. The product was purified by silica column chromatography with a 2:1 hexane:ethyl acetate mobile phase. Norbornene-PEG was synthesized as previously described by reacting norbornenyl NHS ester (1.5 equiv.) and methoxy-PEG amine HCl salt (JenKem Technology USA, 1 equiv.) in the presence of *N,N*-diisopropylethylamine (DIPEA) (Thermofisher, 1.5 equiv.) in DCM for 16 h at room temperature (*70*). The product was precipitated in cold anhydrous ethyl ether, washed 3x, and lyophilized for storage at −20° C. Third-generation Grubbs’ catalyzed was made by reacting 2^nd^ Generation Grubbs Catalyst (MilliporeSigma, 1 equiv.) with 3-bromopyridine (MilliporeSigma, 100 equiv.), followed by precipitation in cold hexane as previously described (*71*).

Diblock bottlebrush polymer was synthesized via ring-opening polymerization of norbornenyl bromide and norbornenyl PEG using 3^rd^ generation Grubbs’ catalyst (*27*). Purified bottlebrush polymer was conjugated with an azide group by substitution reaction with sodium azide (Thermofisher, 100 equiv.) Azide-functionalized bottlebrush polymer (1 equiv.) and DBCO-modified DNA (2.2 equiv.) were reacted in a standard copper-free “click chemistry” reaction for 24 h at 55° C as previously described (*36*). The conjugated pacDNA product was purified by aqueous gel permeation chromatography (GPC) with a 1 M sodium nitrate mobile phase in an Ultrahydrogel 250 angstrom column. DMF-GPC was performed to quantify the molecular weight, PDI, and yield of the bottlebrush polymer. Gel electrophoresis was performed using 2% agarose gel in 1x tris/borate/EDTA (TBE) buffer with a running voltage of 120 V. Gel images were acquired on an Alpha Innotech Fluorochem Q imager. DLS and ζ potential data were recorded on a Malvern Zetasizer Nano-ZSP (Malvern, UK). After DNA conjugation and purification, the pacDNA was desalted using a NAP-25 column (Cytiva) and lyophilized for storage at −20° C.

### Fluorescence in situ hybridization and confocal microscopy

Human fibroblasts were seeded in a 12-well confocal plate and treated with various concentrations of pacDNA or controls for 24 h. Then, cells were fixed in 4% PFA at 4° C followed by permeabilization with 70% ethanol overnight. Pre-hybridization was then performed using a buffer (30% formamide, 2x SSC) for 10 min. Hybridization was performed using a FISH probe 5’-(Cy-3)-C**A**GC**A**GC**A**GCAG**C**AG**C**AG**C**A-3’ (bold=LNA modified, 2 ng/µL) in a buffer containing formamide, SSC, BSA, yeast tRNA, dextran sulfate, and vanadyl ribonucleoside complex (*69*). Cells were stained with DAPI (Thermofisher) for 10 min and washed 3x with PBS. Images were acquired on a Zeiss LSM 880 confocal laser scanning microscope under identical settings (Carl Zeiss Ltd., Cambridge, UK). Foci were counted by eye in ~250 nuclei in each sample group. Confocal imaging of live cells was conducted by a similar procedure, omitting the steps of fixation, permeabilization, and hybridization.

### RNA isolation

HSA-LR20b (HSA^LR^) or FVB/N wildtype mice were euthanized, and muscle samples were collected from quadriceps and flash frozen in liquid nitrogen and stored at −80° C. Individual sections were homogenized in a lysis buffer containing 10% BME on a Beadblaster 24R Microtube Homogenizer (Benchmark scientific, NJ, USA.). Total cellular RNA was extracted using the RNeasy® Fibrous Tissue Mini Kit (Qiagen), and RNA quality and quantity were assessed on a NanoDrop® ND-1000 UV-Vis Spectrophotometer.

### ^89^Zr Radiolabeling to measure biodistribution

For radiolabeling, pacDNA was synthesized with two desferrioxamine (DFO, siderophore-derived chelator) functionalities per molecule, which were used to chelate ^89^Zr-oxalate at a ratio of approximately 500 kBq to 1 μg. The radiochemical labeling yields were monitored by instant radio-thin-layer chromatography. Radio-fast protein liquid chromatography (FPLC) with a size-exclusion column was used to remove unchelated ^89^Zr. To evaluate biodistribution/clearance and perform small animal PET imaging, CD-1 mice were i.v. injected with approximately 500 MBq of pacDNA in 100 μL saline. Mice were anesthetized with inhaled isoflurane and re-anesthetized before euthanasia by cervical dislocation at predetermined time points (24 h, 3 d, 7 d, and 14 d post-injection, n=4/time point; group size based on prior studies). Organs/tissues of interest were collected, weighed, and radio-counted. Standards were prepared and measured along with the samples to calculate the percentage of the injected dose per gram of tissue (%ID/g). Positron-emission tomography (PET) scans were carried out at two of the four time points (24 h and 7 d) prior to the sacrifice of the mice. The microPET images were corrected for attenuation, scatter, normalization, and camera dead time and co-registered with microCT images.

### RNA sequencing

Isolated RNA from quadriceps was sent for total RNA sequencing (Novogene), targeting 40 million reads per sample. After polyA enrichment and quality control, libraries were sequenced on an Illumina PE150 system to obtain paired-end reads of 150 bp. After trimming and alignment, reads were mapped to the mouse genome (mm39), and analysis was performed in RStudio using DESeq2, clusterProfiler, fgsea, gsva, and Rsubread packages. The Benjamini-Hochberg method was applied to account for multiple testing. Adjusted p values (padj < 0.05) were then used to filter differentially expressed genes (DEGs). Gene set enrichment analysis (GSEA) and gene set variation analysis (GSVA) were performed to assess functional gene expression changes after pacDNA-L9 treatment. To identify genes showing recovery patterns after treatment, we calculated recovery scores for each differentially expressed gene between WT and NT conditions. The recovery score was defined as the ratio of the absolute difference between the 42.4 mg/kg pacDNA-L9 treated cohort vs. WT expression to the sum of absolute differences between the 42.4 mg/kg cohort vs. WT and the 42.4 mg/kg cohort vs. NT expression. Lower scores indicate better recovery towards the WT state. The top 25% of genes with the lowest recovery scores were selected for heatmap visualization. Gene expression values were normalized using DESeq2’s size factors and transformed into z-scores for heatmap visualization. Levels of endogenous genes containing short (CUG)_n_ (n=2-25) RNA tracts were examined for potential off-target effects of the pacDNA (Table S1).

Splicing events were calculated using the rMATS statistical model (*72*). Significant splice events were classified as having an FDR < 0.01 and at least 10 junction counts per sample. The composite scaled PSI (cPSI) for individual splicing event x was calculated using the following formula, where PSI(WT) and PSI(NT) are the average PSI values in the wildtype and nontreated cohorts, respectively:

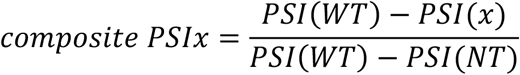

The percent corrected PSI for individual splicing event x was calculated using the following formula, where PSI(WT) and PSI(NT) are the average PSI values in the wildtype and nontreated cohorts, respectively:

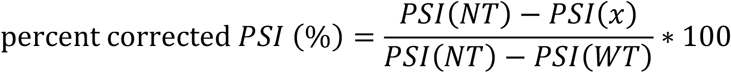

### Behavioral Evaluation of Myotonia and Grip Strength in HSA^LR^ Mice

#### Tail-Pinch–Evoked Myotonia Scoring Assay

To evaluate skeletal muscle myotonia, a blinded behavioral assay was conducted based on a series of tail-pinches. Mice were gently pinched on the lower back, approximately 1 cm cranial to the base of the tail, 15 times per session using blunt forceps. A semi-quantitative scoring system was applied to assess the severity of delayed hindlimb relaxation following each pinch, defined as follows:

- **3** = severe myotonia (bilateral hindlimb extension lasting >1 s)
- **2** = moderate myotonia (0.5–1 s)
- **1** = mild myotonia with quick recovery (<0.5 s)
- **0.5** = unilateral hindlimb extension
- **0** = no myotonic response

Mice were tested in a randomized order to avoid handling bias, and a 20-minute rest was provided after every eight pinches to minimize fatigue. A mean score was calculated from the 15 trials to represent the myotonia severity of each individual mouse.

#### Grip Strength and Body Weight Measurements

Forelimb and hindlimb grip strength was measured using a Grip Strength Meter (Bioseb) at baseline and selected time points throughout the treatment period. Four independent readings were taken per mouse per session, and the average value was recorded. Results were compared with age- and sex-matched untreated HSA^LR^ and FVB/N wild-type control mice. Body weight was measured weekly using a calibrated digital scale.

#### Anti-PEG immune response

HSA^LR^ mice were administered PBS or pacDNA-L9 (10.6 mg/kg) weekly. As a positive control for PEG immunogenicity, a separate group of HSA^LR^ mice received 4 mg/kg of polyethylene glycol–keyhole limpet hemocyanin (PEG-KLH), a known immunogenic PEG conjugate. Plasma (100 µL) was collected one and three weeks after the final dose. Anti-PEG IgG and IgM concentrations were measured using ELISA kits (Cat #PEGG-1 and #PEGM-1, Life Diagnostics, Inc.) according to the manufacturer’s protocol.

#### Multiplex analysis of cytokines and blood markers

HSA^LR^ mice were treated with pacDNA-L9, free L9 ASO, PBS, or lipopolysaccharide (LPS, 2 mg/kg) positive control. Plasma was collected 4 h after the final injection and multiplexing analysis of cytokines was performed using the Luminex™ 200 system (Luminex, Austin, TX, USA) by Eve Technologies Corp. (Calgary, Alberta). The 32-plex panel included Eotaxin, GM-CSF, IL-1α, IL-1β, IL-2, IL-3, IL-4, IL-5, IL-6, IL-10, IL-12(p70), IL-13, IL-15, KC, LIX, MCP-1, and TNFα. The assay sensitivities of these markers range from 0.3–30.6 pg/mL. Individual analyte sensitivity values are available in the MilliporeSigma MILLIPLEX® MAP protocol. Serum chemistry biomarkers (Albumin, ALT, AST, ALP, blood urea nitrogen, calcium, creatinine kinase, creatinine, phosphorous, total protein, and total bilirubin) were quantified by Idexx Bioanalytics (Columbia, MO) according to company protocols.

#### Statistical analysis

For two-group and multigroup comparisons, we used unpaired two-tailed t tests or multigroup analysis of variance (ANOVA) (GraphPad Prism 10.4.1 software, GraphPad, Inc.). We used the F test to compare variances between the two groups. P-values denoted by * symbols were calculated in Prism. Data are presented as the mean ± s.e.m. or mean ± s.d., with n values noted in all figure captions. A two-tailed nonparametric *t*-test was applied to compare two groups that have a statistically significant difference in variance (Foci counting, myotonia, body weight). One-way ANOVA followed by Tukey’s multiple comparison test was applied for composite PSI analysis of the DM1 splicing panel and grip strength functional study.

## Supporting information

Supplementary Materials, includes: Figs. S1 to S7, Table S1, Abbreviations.

## Acknowledgments

We are grateful to the Northeastern University-Institutional Animal Care and Use Committee for supporting our animal studies. We thank Dr. Guoxin Rong from the Institute for Chemical Imaging of Living System at Northeastern University for assistance with confocal microscopy. We appreciate Prof. Sara H Rouhanifard for guidance and resources for performing FISH experiments.

## Funding

This publication was made possible by the

National Science Foundation DMR award number 2004947 (KZ)

National Institute of General Medical Sciences 1R01GM121612 (KZ)

Department of Defense CDMRP award number MD230034 (KZ)

## Author contributions

Conceptualization: KZ, YL, CO

Methodology: YL, CO

Investigation: YL, CO, KZ

Bioinformatic Analysis: JW, CO, YL

Biodistribution study: YW, GSH, YL

Confocal experiments: YL, RC, HZ, KN, CO

Supervision: KZ

Writing—original draft: CO, YL, KZ

Writing—review & editing: CO, YL, KZ

All other authors contributed to material synthesis. All authors have given approval to the final version of the manuscript.

## Data and materials availability

All data are available in the main text or the supplementary materials.

