## Supplementary Materials, includes: Figs. S1 to S7, Table S1, Abbreviations. for "Bottlebrush Polymer Conjugates for Enhanced Antisense Oligonucleotide Therapy in Myotonic Dystrophy Type 1"

**This PDF file includes:**

Figs. S1 to S7
Table S1
Abbreviations

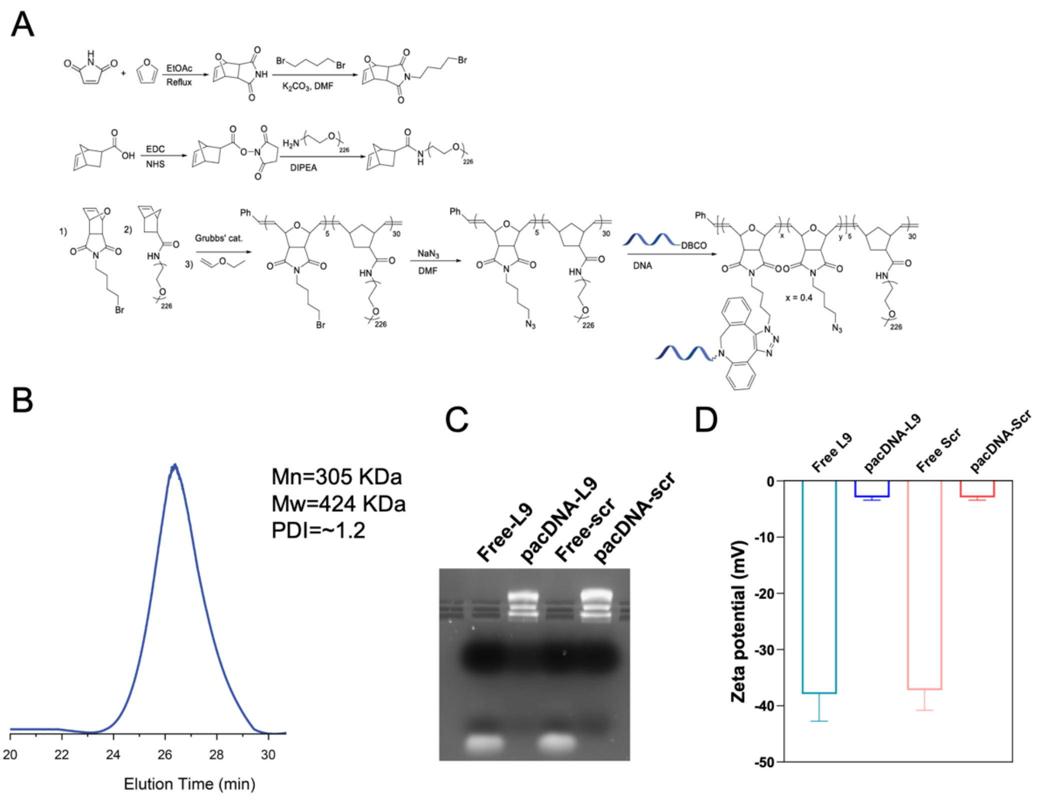


**Fig. S1. (A)** Synthesis scheme of the pacDNA conjugate. DBCO-functionalized ASO shown in blue **(B)** Aqueous gel permeation chromatograph of the pacDNA-L9 conjugate. Molecular weight (Mn and Mw) and polydispersity (PDI) were approximated by DMF-GPC. **(C)** Agarose gel (2%) electrophoresis separation of free L9 ASO (lane 1), pacDNA-L9 (lane 2), free L9-Scr ASO (lane 3), and pacDNA-Scr (lane 4). **(D)** Zeta potential (ζ) measurements of the pacDNA-L9, pacDNA-Scr, free L9 ASO, and L9-Scr ASO in Nanopure water. Error bars indicate ±s.d.

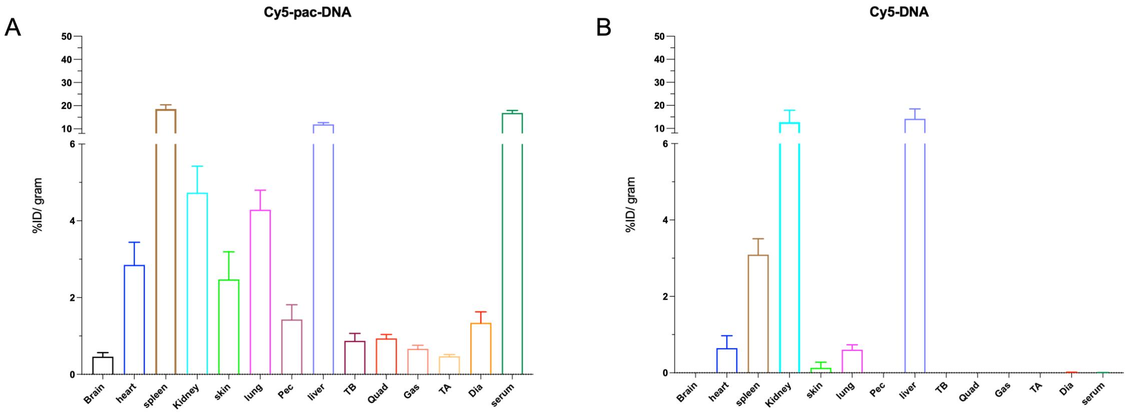


**Fig. S2. Biodistribution of Cy5-labeled pacDNA and free ASO. (A, B)** Ex vivo biodistribution of Cy5-pacDNA and Cy5-DNA in various tissues and organs measured over 3 days (n=3) in $HSA^{LR}$ mice. The y axis shows the percent of injected dose per gram of tissue (%ID/g). Quad-quadriceps; Gas-gastrocnemius; TA-tibialis anterior; Pec–pectoralis; Dia–diaphragm.

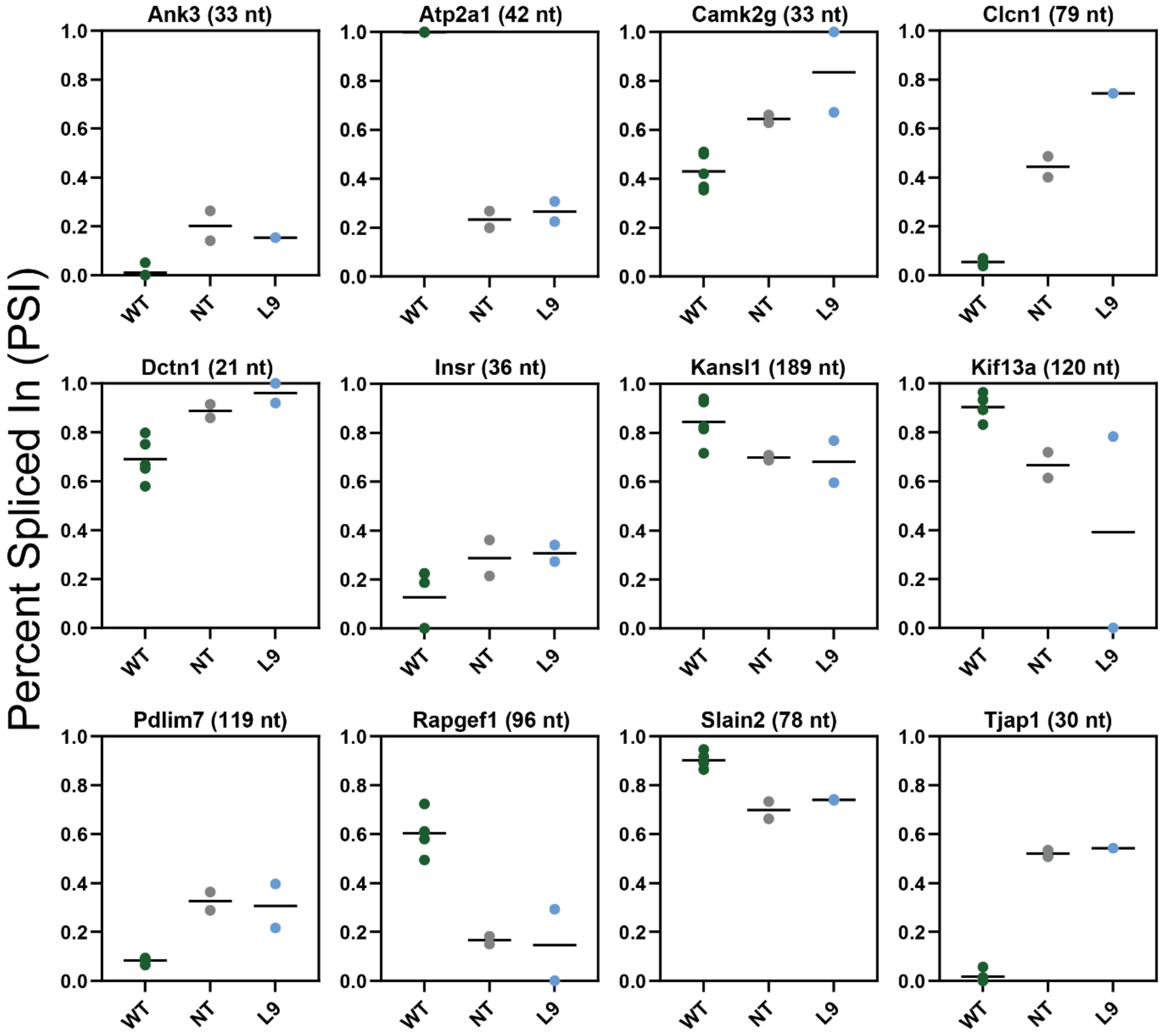


**Fig. S3.** Percent spliced-in (PSI) graphs for 12 key DM1-associated splice events in wildtype (n=5), nontreated HSA$^{LR}$ (n=2) and free L9 ASO-treated HSA$^{LR}$ mice (5.3 mg/kg, n=2) in quadriceps two weeks post-injection.

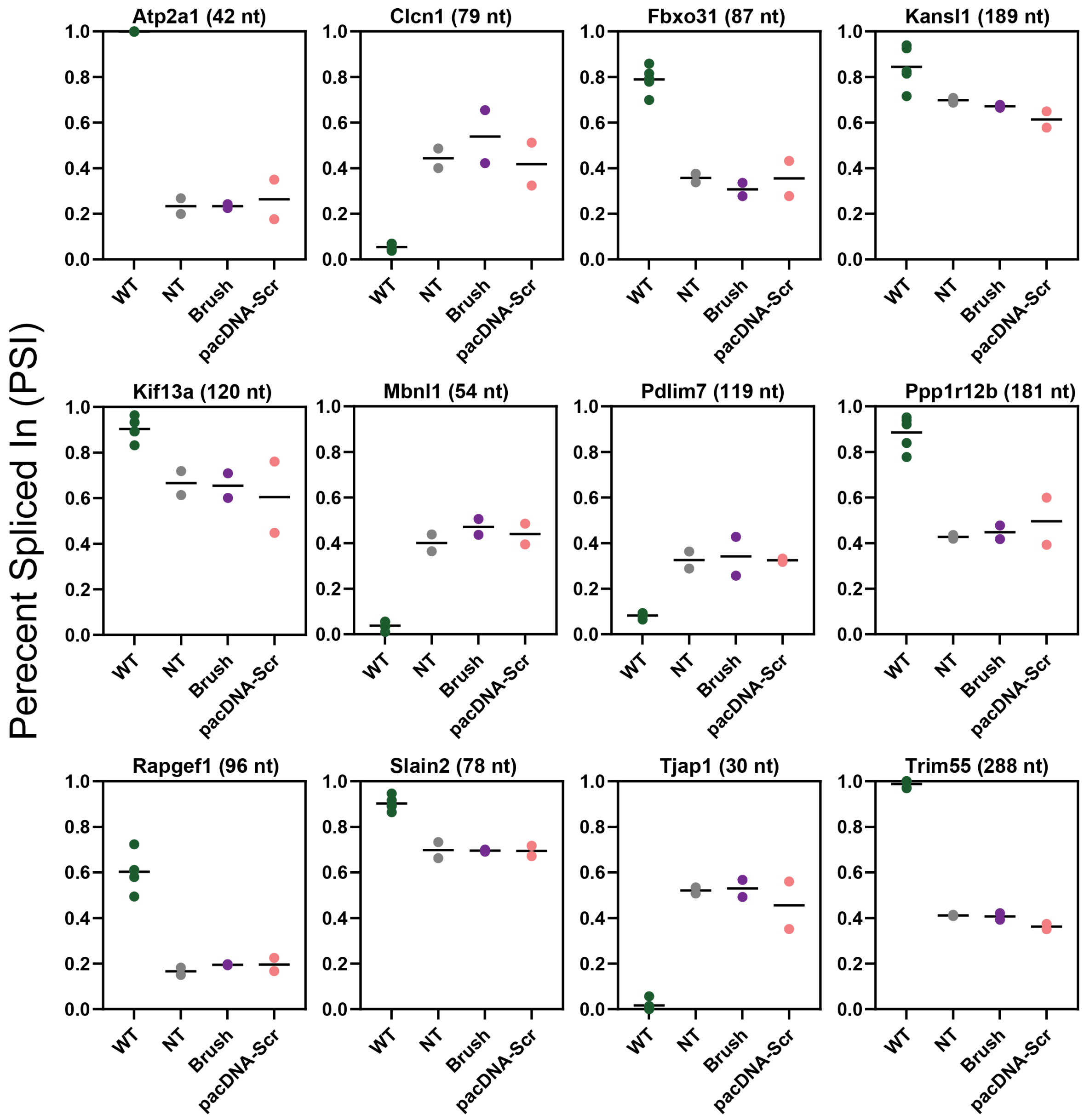


**Fig. S4.** Percent spliced in (PSI) of 12 key DM1-associated splice events in wildtype (n=5), nontreated HSA$^{LR}$ (n=2), brush (equimolar to pacDNA-L9, n=2) and pacDNA-Scr (5.3 mg/kg, n=2) treated HSA$^{LR}$ quadriceps, two weeks post-injection.

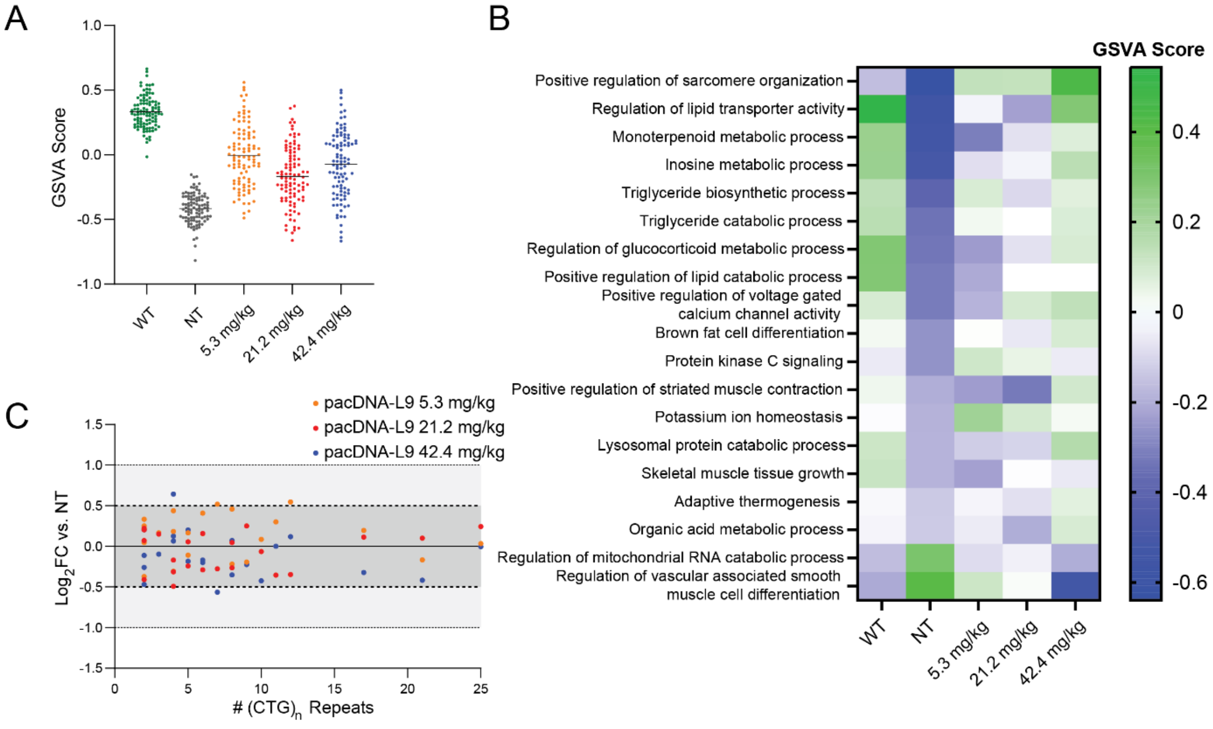


**Fig. S5. (A)** Gene set variation analysis (GSVA) scores of the 100 lowest score (most downregulated) GO biological processes in nontreated HSA$^{LR}$ samples compared to wildtype across all treatment groups. **(B)** Select GO biological process GSVA scores visualized by heatmap across all treatment groups. **(C)** Expression levels of endogenous genes containing short $(CUG)_n$ (n=2-25) RNA tracts in pacDNA-L9 treated mice vs. NT (Table S1).

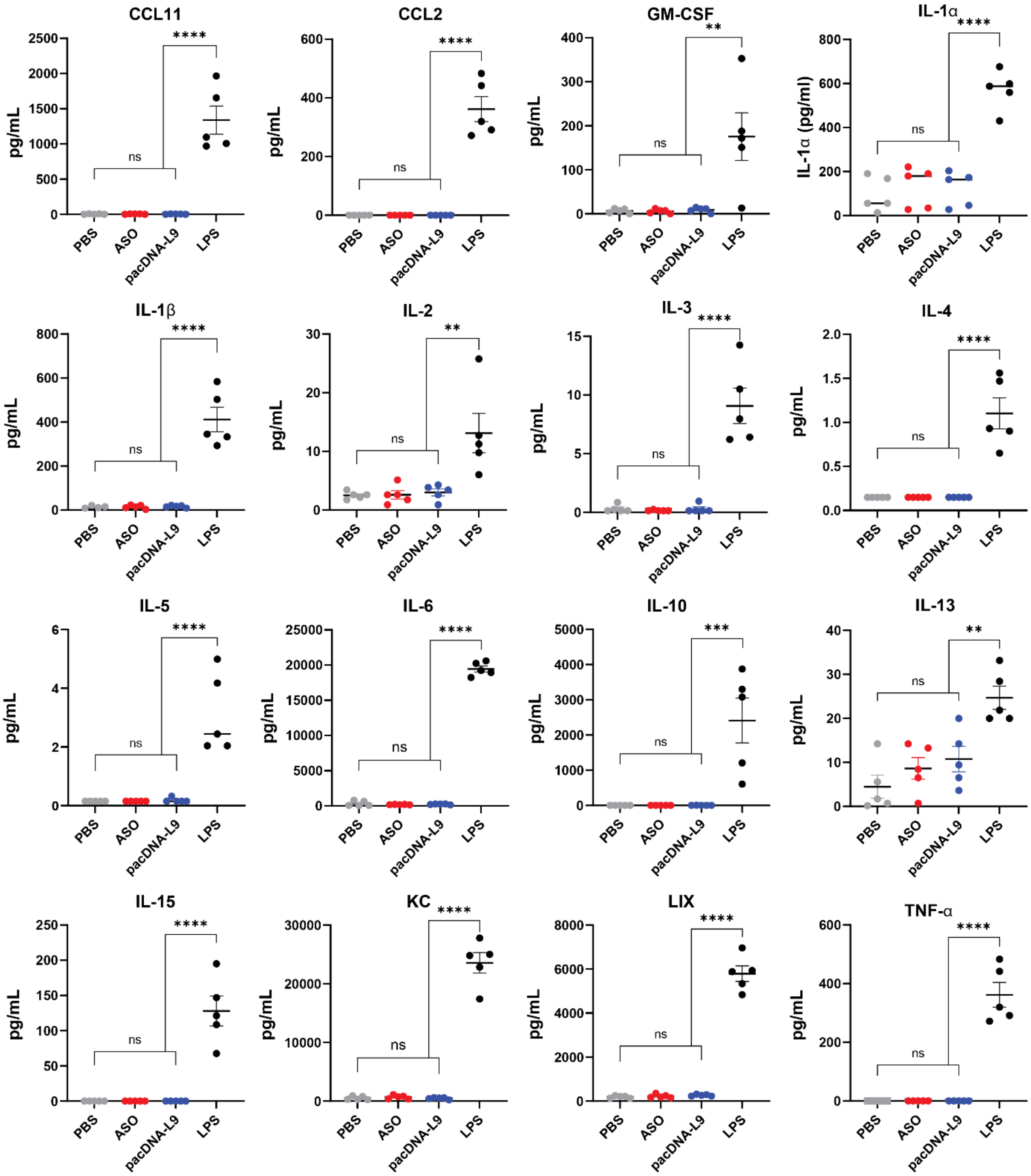


**Fig. S6.** Cytokine and chemokine levels in HSA$^{LR}$ mice serum 4 h after i.v. injection of PBS, free L9-ASO, pacDNA-L9 (10.6 mg/kg), or LPS (2 mg/kg) (n=5 per group) (****p<0.0001, ***p<0.001, **p<0.01, one-way ANOVA). Error bars indicate ±s.e.m.

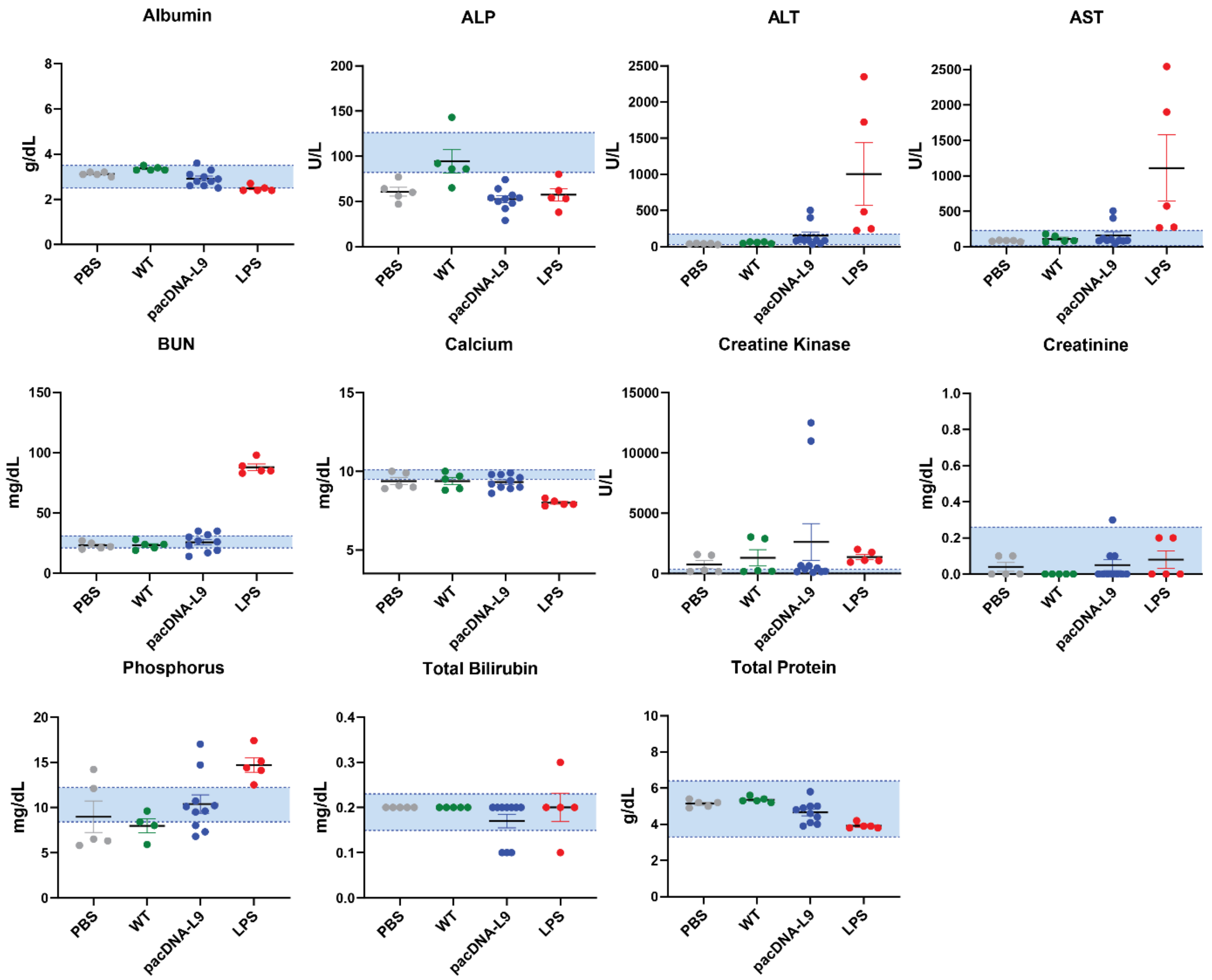


Fig. S7. Liver function and renal function biomarker levels in HSA$^{LR}$ mice serum collected after i.v. injection of PBS (n=5), pacDNA-L9 (10.6 mg/kg) (n=10), or LPS (2 mg/kg) (n=5), or in FVB/N wildtype (WT) mice (n=5). The blue shaded area represents the healthy reference range in FVB/N mice. Error bars indicate ±s.e.m.

| Gene | $(CTG)_n$/ *$(CAG)_n$ | $Log_2FC$ pacDNA-L9 5.3mg/kg vs. NT | $Log_2FC$ pacDNA-L9 21.2mg/kg vs. NT | $Log_2FC$ pacDNA-L9 42.4mg/kg vs. NT |
|---|---|---|---|---|
| Papss2 | 2 | 0.251 | 0.123 | 0.121 |
| Bpgm | 2 | -0.255 | -0.29 | 0.006 |
| Dmpk | 2 | 0.011 | 0.189 | -0.291 |
| Tcf4 | 2 | 0.072 | -0.106 | -0.646 |
| Ltbp3 | 3 | 0.196 | 0.184 | -0.064 |
| Rpl14 | 4 | -0.198 | -0.202 | 0.176 |
| Lrp8 | 4 | 0.166 | -0.757 | 0.352 |
| Bri3bp | 4 | 0.328 | -0.022 | 0.27 |
| Map3k4 | 5 | -0.118 | 0.052 | -0.192 |
| Notch4 | 5 | -0.089 | -0.491 | -0.049 |
| Ptbp1 | 6 | 0.25 | -0.445 | -0.355 |
| Sdc3 | 6 | 0.084 | 0.089 | -0.24 |
| Armcx6 | 7 | 0.642 | -0.145 | -0.438 |
| Mllt3 | 8 | -0.18 | -0.222 | -0.313 |
| Tacc1 | 8 | 0.271 | -0.137 | -0.114 |
| Txlnb | 9 | -0.231 | 0.218 | -0.261 |
| Tnfrsf22 | 10 | 0.32 | 0.174 | -0.189 |
| Pcolce | 12 | 0.428 | -0.462 | 0 |
| Fgd4 | 21 | -0.315 | -0.043 | -0.564 |
| Mapkap1 | 25 | 0.089 | 0.302 | 0.053 |
| Nr3c1* | 17 | 0.114 | 0.033 | -0.401 |
| Dap* | 11 | 0.045 | 0.039 | 0.178 |

**Table S1**. Expression levels of endogenous genes containing short (2-25) CTG repeats relative to non-treated (NT) $HSA^{LR}$ mice.

**Abbreviations:**

3'-UTR: Three prime untranslated region
ASO: Antisense oligonucleotide
CLCN1: Chloride voltage-gated channel 1
CPG: Controlled-pore glass
$CUG^{exp}$: CUG-repeat expanded RNA
DBCO: Dibenzocyclooctyne
DFO: Desferrioxamine
DLS: Dynamic light scattering
DM1: Myotonic dystrophy type 1
DM2: Myotonic dystrophy type 2
DMD: Duchenne muscular dystrophy
DMPK: DM1 Protein Kinase
DMT: Dimethoxytrityl
DIPEA: *N*,*N*-diisopropylethylamine
EDCI: 1-ethyl-3-(3-dimethylaminopropyl)carbodiimide
ELISA: Enzyme-linked immunosorbent assay
FDR: False discovery rate
FISH: Fluorescence *in situ* hybridization
FSHD: Facioscapulohumeral muscular dystrophy
GPC: Gel permeation chromatography
GSEA: Gene set enrichment analysis
GSVA: Gene set variation analysis
I.V.: Intravenous
KLH: Keyhole limpet haemocyanin
LNA: Locked nucleic acid
LPS: Lipopolysaccharide
MBNL1: Muscleblind-like one
pacDNA: Polymer-assisted compaction of DNA
PEG: Polyethylene glycol
PMO: Phosphorodiamidate morpholino oligomer
PS: Phosphorothioate
PSI: Percent spliced in
rMATS: R multivariate analysis of transcript splicing
RP-HPLC: Reversed-phase high-performance liquid chromatography
SMA: Spinal muscular atrophy
$^{89}Zr$: Zirconium-89
